# Assessing Target of Rapamycin (TOR) activity in the diatom *Phaeodactylum tricornutum* using commercially available materials

**DOI:** 10.1101/2024.03.28.587301

**Authors:** Yoshinori Tsuji, Takumi Ishikawa

## Abstract

Target of rapamycin (TOR) is a conserved protein kinase that regulates the balance between catabolic and anabolic processes in response to nutrient availability. Although the central role of TOR kinase in nutrient stress responses is well-recognized, little is known about the molecular basis of TOR signaling in ecologically important secondary algae with plastids of red algal origin, such as diatoms, as assessing *in vivo* TOR kinase activity is a difficult task. To assess TOR kinase activity, the phosphorylation status of downstream components, such as ribosomal protein S6 (RPS6), must be measured. Unlike for model organisms, an antibody that detects phosphorylated (P-) RPS6 in diatoms is not commercially available. Therefore, we developed a convenient method in which P-RPS6 and non-P-RPS6 were detected via Phos-tag affinity electrophoresis and immunoblotting with a commercial antibody that cross-reacts with RPS6 (both P- and non-P-RPS6) in the diatom, *Phaeodactylum tricornutum*. Using this Phos-tag-based method, we observed a dose-dependent decrease in the P-RPS6/total RPS6 ratio in *P. tricornutum* cells treated with the TOR kinase inhibitor, AZD-8055. We also observed a reduction in the P-RPS6/total RPS6 ratio during the nitrogen-deficient culture of *P. tricornutum*, which indicated the inactivation of TOR kinase in response to nitrogen deficiency. Finally, we demonstrated the potential application of the Phos-tag-based method to other ecologically, evolutionarily, and industrially important secondary algae, such as *Nannochloropsis oceanica* (Stramenopiles), the haptophyte *Tisochrysis lutea*, and *Euglena gracilis* (Euglenid). As all experimental materials are commercially available, the Phos-tag-based method can be used to promote studies on TOR in diverse algae in different contexts.

## 1. Introduction

Target of Rapamycin (TOR) is a central protein kinase that coordinates eukaryotic growth and development. Since its discovery, TOR has been extensively studied in non-photosynthetic eukaryotes, such as yeast and mammals [1]. These organisms have two distinct TOR complexes with different functions, TOR complex 1 (TORC1) and TOR complex 2 (TORC2), whereas plants and algae only have TORC1 [2]. TORC1 is activated when nutrients are sufficient to promote anabolic processes and cell division, whereas nutrient depletion induces opposite responses. In mammalian cells, nutrient sensing by TORC1 is mediated by the Ragulator-Rag complex located on the lysosomal surface [1]. The Ragulator-Rag complex contains two small GTPases (Ras-related GTP-binding proteins, Rag A/B and C/D) that act as molecular switches to modulate the localization of TORC1. Increases in cellular amino acid levels alter the nucleotide-bound state of Rag, resulting in the relocation of TORC1 onto the lysosomal surface, where TORC1 is allosterically activated by another small GTPase, Ras Homolog Enriched in Brain (Rheb).

In contrast to mammals, whose primary nutrients are ingested as organic compounds, plants and algae assimilate inorganic nutrients through photosynthesis. This notable difference in nutrient acquisition raises the fundamental question of how TORC1 senses cellular nutrient levels in phototrophs. According to a database analysis, plants lack mammalian-type signaling components, such as Ragulator, Rag, and Rheb [2]. Recent studies have shown that a plant-specific small GTPase, Rho-related protein from plant 2 (Rop2), transmits nutrient signals to TORC1 in *Arabidopsis thaliana* [3]. Stimuli from various nitrogen sources convert Rop2 into its GTP-bound active form, leading to the activation of TORC1 (Liu et al., 2021). Notably, Rop2 also activates TORC1 via auxins [4], suggesting that ROP2 is a central hub that rectifies diverse upstream signals. Recent studies have suggested that both inorganic and organic nitrogen can activate TORC1 in *A. thaliana* [3,5,6]; however, the detailed mechanism has not been elucidated. The inclusion of whole eukaryotic phototrophs suggests further complexity in TORC1 signaling, which may have arisen from multiple endosymbiotic evolutions [2]. Among the diverse algal lineages, TORC1 has been mainly studied in Archaeplastida, whose plastid is derived from a cyanobacterial endosymbiont, such as the green alga *Chlamydomonas reinhardtii* (*Chlamydomonas* hereafter) and the red alga *Cyanidioschyzon merolae* [7]. In *Chlamydomonas*, TORC1 activity is regulated by the availability of nitrogen [8], inorganic phosphate (Pi) [9], and CO_2_ [10]. However, *Chlamydomonas* lacks both plant- and mammalian-type signaling components, such as Rop2, Rag, and Ragulator [2,11], suggesting that green algae might have evolved a unique system in the regulation of TORC1.

Unlike Archaeplastida, ecologically important algal groups, such as diatoms (Stramenopiles), have evolved via secondary endosymbiosis, in which a red algal endosymbiont is engulfed as a plastid. Due to their presumed serial endosymbiosis with green and red algae, diatoms have a chimeric genome consisting of genes from diverse sources [12]. Diatoms account for 20% of the net primary production on Earth, driving the global carbon cycle and aquatic ecosystems. Owing to the high capacity of these diatoms to accumulate triacylglycerols (TAGs) under nutrient starvation conditions, they have attracted attention as a potential source of renewable energy [13]. In the pennate diatom, *Phaeodactylum tricornutum*, chemical inhibition of TORC1 induces TAG accumulation and growth retardation in a manner similar to that in nitrogen-deprived cells [14]. However, only physiological analyses have been conducted using TOR inhibitors in diatoms, and the molecular basis of TORC1 signaling has not been unexplored [2,15]. Interestingly, some mammalian-type signaling components appear to be conserved in diatoms [2], but their involvement in TORC1 regulation has not been investigated yet.

A major constraint that limits studies on algal TORC1, especially in secondary lineages, is the lack of a method for assessing *in vivo* TORC1 activity. In model organisms, such as mammals, the phosphorylation status of downstream effectors, such as ribosomal S6 kinase (S6K) and ribosomal protein S6 (RPS6), is commonly used as a readout of *in vivo* TORC1 activity [16,17]. S6K is phosphorylated and activated in a TORC1-dependent manner, leading to phosphorylation of RPS6 by S6K. Owing to the high conservation of the TORC1-S6K-RPS6 axis, these downstream effectors are widely used as indicators of TORC1 activity in diverse organisms, such as yeast, mammals, and plants [16–18]. Commercial antibodies that react with phosphorylated (P-) S6K and/or P-RPS6 are available for mammals and plants, enabling easy monitoring of *in vivo* TORC1 activity.

In contrast to model organisms, such antibodies are not commercially available for algae, making it difficult to assess algal TORC1 activity. Several efforts have been made to detect P-S6K or P-RPS6 in the model alga, *Chlamydomonas*, using commercial antibodies. Early attempts to use antibodies against mammalian P-S6K were unsuccessful [19,20]. More recently, Upadhyaya et al. (2020) succeeded in detecting P-S6K in *Chlamydomonas* using a commercial anti-mammalian P-S6K; however, the detected mass (35 kDa) was not consistent with the theoretical mass of CrRPS6 (98.8 kDa), thereby warranting further validation. In addition to trials with commercial antibodies, Couso et al. (2020) opted to generate specific antibodies to detect P-CrRPS6 (Ser245) and total CrRPS6, establishing a reliable method to quantify P-RPS6/total RPS6 ratio as a readout of TORC1 activity. Although generating antibodies is an authentic method, the same strategy cannot be easily applied to other algae because the regulatory phosphosites tend to vary in different organisms, reinforcing costly processes, such as identification of phosphosites and subsequent generation of antibodies against phosphopeptides. Overall, methods for monitoring TORC1 activity have recently become available in the model green alga, but efforts still need to be made in ecologically and industrially important secondary lineages.

Diatoms play central roles in the supply of organic carbon to oceanic ecosystems. Although TORC1 in diatoms is likely to be the central regulator regulating growth and primary production in response to fluctuating nutrient availability, the lack of methods for monitoring TORC1 activity has hindered molecular dissection of TORC1 signaling. To overcome this limitation and facilitate TORC1 studies in secondary algae, we aimed to establish a convenient method for monitoring diatom TORC1 activity using commercially available materials.

## 2. Materials and Methods

### 2.1. Strains and maintenance culture

*Chlamydomonas reinhardtii* (strain CC-5325) was obtained from Chlamydomonas Resource Center (https://www.chlamycollection.org) and photomixotrophically grown in Tris-Acetate-Pi medium (TAP) at 25 °C. The pennate diatom, *Phaeodactylum tricornutum* (CCAP1055/1, also known as Pt1), was obtained from the National Institute for Environmental Studies, Japan (deposited as NIES-4392) and cultured at 20 °C in artificial seawater Marine Art SF-1 (Osaka Yakken, Osaka, Japan) enriched with 0.1 mM sodium metasilicate and modified Erd-Schreiber medium (ESM) in which soil extract was replaced by 10 nM sodium selenite (MA-ESM medium Danbara & Shiraiwa, 1999). Salinity was set at 30‰ using a refractometer (MASTER-S/MillM; ATAGO, Tokyo, Japan). *Nannochloropsis oceanica* (NIES-2145) and *Tisochrysis lutea* (kind gift from Dr. Iwane Suzuki, University of Tsukuba) were grown under the same conditions as *P. tricornutum*. *Euglena gracilis* (NIES-48) was grown photoautotrophically at 25 °C in AF6 medium buffered with Tris-HCl (pH7.0).

### 2.2 Inhibitor treatments

*Chlamydomonas* was inoculated in 400 mL TAP in a 1 L flask at an initial density of OD_730_ = 0.08, and cultured under orbital shaking (100 rpm) with constant illumination (50 μmol photons m^−2^ s^−1^) at 25 °C. When the OD_730_ reached 0.4, the bulk culture was divided into small cultures (50 mL each in 200 mL flasks), which were administered different concentrations of inhibitors or dimethyl sulfoxide (DMSO, as a control). After 30 min, the cells were harvested via centrifugation (2,350 ×*g*, 2 min, 4 °C), washed with ice-cold extraction buffer (composition is described in the next section), and frozen in liquid N_2_. AZD-8055 (Cat. No. CS-0067; ChemScene, NJ, USA) and rapamycin (Cat. No., R-5000; LC Laboratories, Inc., MA, USA) were used as the TOR inhibitors.

To treat *P. tricornutum* with the inhibitor, cells were precultured in 60 mL MA-ESM in a 200 mL flask with continuous illumination (50 μmol photons m^−2^ s^−1^) and orbital shaking (80 rpm). When the OD_730_ reached 0.2-0.3, the culture was inoculated into 50 mL MA-ESM at an OD_730_ = 0.07 (test tube, φ30 mm) and bubbled with air under constant illumination (80 μmol photons m^−2^ s^−1^). When the OD_730_ reached 0.4-0.5 after 2 days, AZD-8055 or DMSO was added and the cells were further cultured for 2 h. The cells were harvested via centrifugation (2,350 ×*g*, 2 min, 4 °C), washed using 500 μL of ice-cold AP-buffer (composition described in the next section), and stored at −80 °C until use.

### 2.3. Nitrogen-deficient culture of *P. tricornutum*

Cells grown in normal MA-ESM medium containing nitrate (OD_730_ = 0.5-0.6) were harvested via centrifugation (1700 ×*g*), washed twice with 10 mL of nitrate-free MA-ESM, and resuspended in nitrate-free MA-ESM at an OD_730_ = 0.5. To eliminate the effects of centrifugation and medium exchange, the cells were allowed to acclimate to culture conditions (illumination at 80 μmol photons m^−2^ s^−1^ with air bubbling) for 30 min, and then initial sampled (0 h). For the resupply experiments, sodium nitrate (50 mM) or ammonium chloride (50 mM) dissolved in 10 mM Tris-HCl (pH 8.0) was added to the nitrogen-deficient cultures to a final concentration of 500 μM. As a control, 10 mM Tris-HCl (pH8.0) without nitrogen was added.

### 2.4. Preparation of the crude extract for SDS-PAGE

All steps were performed at 4 °C or on ice, unless otherwise mentioned. For *Chlamydomonas*, the cell pellet derived from 50 ml of culture was suspended in 500 μl of the extraction buffer [50 mM Hepes-NaOH (pH7.5), 10 mM NaCl, and cOmplete Mini EDTA-free protease inhibitor cocktail (Cat# 4693159001, Sigma Aldrich, Japan)]. The cells were disrupted using a handy sonicator (Model UR-21P; Tomy Seiko Co., Ltd., Tokyo, Japan) at an intensity of 8 with repeated 3-s pulses at 3-s intervals for 2 min. Centrifugation (20,000 ×*g*, 20 min) and discarding of the pellet were repeated three times to obtain soluble proteins as a crude extract. After the addition of ×6 Laemmli sample buffer, the crude extracts were heated at 56 °C for 5 min and applied to the PAGE gel. For *P. tricornutum*, the cell pellet corresponds to 20 mL of culture suspended in 300 μl of the extraction buffe and disrupted using a sonicator. Soluble proteins were obtained after three rounds of centrifugation (20,000 ×*g*, 20 min), and the supernatant was obtained as the crude extract. Protein concentrations were determined using a Bradford assay kit with bovine serum albumin as a standard (Nacalai Tesque, Kyoto, Japan).

### 2.5. CHCl_3_/MeOH precipitation

In the Phos-tag PAGE analysis with *P. tricornutum* samples, we could not detect signals of P-forms of PtRPS6 using 5 μg of the crude proteins, presumably due to the dispersion of signals into multiple P-forms. Therefore, we applied 20 μg lane^−1^ to detect signals of P-forms by concentrating the samples via CHCl_3_/MeOH precipitation (Wessel & Flügge, 1984). Briefly, 100 μL of the crude extract containing 100 μg of proteins was precipitated and dissolved in 50 μL sample buffer. Thereafter, 10 μL (corresponds to 20 μg of proteins) was dispensed into each lane.

### 2.6. Phosphatase treatment

To determine the band representing non-P-RPS6, crude extracts were treated with the λ-protein phosphatase (λ-PPase; New England BioLabs, Cat# P0753S) before application on the gel. For λ-PPase treatment, the crude extract was diluted to 1 mg proteins mL^−1^ using the extraction buffer. After addition of 1% (v/v) Brij35, 200 mM DTT, and 10 mM MnCl_2_ at final concentrations of 0.01%, 2 mM, and 1 mM, respectively, the crude extract containing 100 μg (in 100 μL volume) of proteins was treated with 400 units of λ-PPase (#P0753; New England Biolabs) for 60 min at 30 °C. After the reaction, *Chlamydomonas* samples were denatured via mixing with ×6 Laemmli sample buffer and heating. For *P. tricornutum*, samples were subjected to CHCl_3_/MeOH precipitation as described in the above section prior to the Phos-tag PAGE analysis.

### 2.7. SDS-PAGE with and without Phos-tag

We employed Zn^2+^-Phos-tag SDS-PAGE with a neutral Bis-Tris gel instead of the Mn^2+^-Phos-tag system owing to the superior mobility shift of phosphorylated proteins with the neutral Zn^2+^-Phos-tag [22]. For Zn^2+^-Phos-tag SDS-PAGE, a gel containing 8% acrylamide was supplemented with 25 μM of Phos-tag and 50 μM of ZnCl_2_ (See supplemental material 1 and 2 for details). For electrophoresis without Phos-tag and ZnCl_2_, 12% acrylamide gel was used. SDS-PAGE was performed using a CompactPAGE system (WSE-1025; ATTO, Tokyo, Japan) and the standard settings (20 mA per gel for 45-55 min). Of note, the addition of sodium bisulfide to the running buffer of Phos-tag PAGE is essential to prevent band distortion. For normal SDS-PAGE, crude extract containing 5 μg of proteins was dispensed into each lane for samples from *Chlamydomonas* and *P. tricornutum*. For Phos-tag PAGE, 5 μg and 20 μg of proteins were dispensed into each lane for samples from *Chlamydomonas* and *P. tricornutum*, respectively.

### 2.8. Immunoblotting

After Phos-tag PAGE, the Zn^2+^-Phos-tag complex in the gel was deactivated by washing the gel five times (10 min each) with chelating buffer (10 mM EDTA, 40 mM Tris, 242 mM glycine, and 0.05% SDS). This procedure was omitted in the normal gel without Phos-tag. The gel was equilibrated with blotting buffer for 10 min and the proteins were blotted onto a PVDF membrane (FluoroTrans W Membrane, PALL Life Science) via semidry blotting at 2 mA cm^−2^ for 1.5 h (Phos-tag gel) or 1.0 h (normal gel) using Bjerrum Schafer-Nielsen buffer supplemented with 0.05% SDS, 5 mM EDTA, and 5% (v/v) MeOH (Table S1). For the blotting of proteins on a normal gel (without the Phos-tag), SDS was reduced to 0.02% (Table S1). After blotting, the membranes were blocked with 3% skim milk in phosphate buffered saline (PBS), incubated with the primary antibodies, anti-mammalian RPS6 (Cat# 2317S; Cell Signaling Technology, Danvers, MA, USA) against *Chlamydomonas* and anti-mammalian RPS6 (Cat#ab225676; Abcam, Cambridge, UK) against *P. tricornutum*, for 1.5 h, and then incubated with goat anti-rabbit IgG conjugated to horseradish peroxidase (HRP) (Cat# 65-6120; Invitrogen) and anti-mouse IgG HRP conjugate (W4021; Promega, Madison, WI, USA) as secondary antibodies for 1.5 to 2.0 h at room temperature with gentle shaking. For the secondary antibody reaction, 0.5% skim milk was added to PBS containing 0.05% Tween 20 (PBST), which effectively suppressed nonspecific signals. Detailed information is provided in the supplemental material 3. The signal was developed using a chemiluminescence reagent (Immobilon Forte #WBLUC0500; Merck, Keniworth, NJ, USA) and detected using a high-sensitivity imaging system (LAS-4000, GE healthcare, Chicago, IL, USA).

### 2.9. Growth monitoring on the agar medium

TAP containing 1.5% agar or MA-ESM containing 1.2% agar was used to monitor the growth of *Chlamydomonas* or *P. tricornutum*, respectively. Cells in the logarithmic growth phase were adjusted to OD_730_ = 0.3 and dispensed onto the solid medium following five-fold serial dilutions (×1, ×1/5, and ×1/25). *Chlamydomonas* was grown at 25 °C under 50 μmol photons m^−2^ s^−1^ while *P. tricornutum* was grown at 20 °C under 50 μmol photons m^−2^ s^−1^.

## 3. Results

### 3.1. Separation of P-RPS6 and non-P-RPS6 in the model alga, *Chlamydomonas*

As carried out in model organisms, we used the conserved TORC1-S6K-RPS6 axis as an indicator to monitor TORC1 activity. As the downstream components of the cascade are abundant and easy to detect, we selected RPS6 as an indicator. To avoid costly and time-consuming steps, such as identifying phosphorylated residues and generating anti-phosphopeptides, we used Phos-tag affinity electrophoresis (Phos-tag PAGE), in which the Phos-tag captures P-forms, causing mobility shifts relative to the non-P-form.

As Couso et al. (2020) previously demonstrated that Ser245 on CrRPS6 underwent TORC1-dependent phosphorylation using anti-P-CrRPS6, we tested Phos-tag-based separation of RPS6 in the model green alga, *Chlamydomonas* (CrRPS6). To detect CrRPS6 in the Phos-tag and normal gels, we selected a mouse monoclonal antibody generated against human RPS6 (Cat# 2317S; Cell Signaling Technology) because this antibody was previously used in *A. thaliana* (Dobrenel et al., 2016). When the crude extract of *Chlamydomonas* was resolved by normal SDS-PAGE (without the Phos-tag), a single band (29.1 kDa) was detected, aligning with the mass calculated from the sequence (28.4 kDa, Phytozome ID Cre09.g400650; Fig. S1).

To determine the phosphorylation status of CrRPS6, cells were treated with various concentrations of TOR inhibitors and their crude extracts were analyzed via Phos-tag PAGE and immunoblotting. In cells that were not treated with an inhibitor (0 min or Ctrl lanes in Fig. 1A), at least 4 different bands were detected (P0–P3 in Fig. 1A), with P0 identified as the non-P form by phosphatase treatment prior to Phos-tag PAGE (Fig. S2). P1 was identified as the major P-form, accounting for approximately 40% of the total CrRPS6, whereas P2 and P3 were the minor forms, each accounting for approximately 10% of the total (Fig. 1B). Cells treated with the ATP-competitive TOR inhibitor, AZD-8055, exhibited a dose-dependent decrease in the P-CrRPS6/total CrRPS6 ratio from 61% in the control to 15% (Fig. 1A and B). Treatment with the allosteric TORC1 inhibitor, rapamycin (500 nM), also decreased the P-RPS6/total RPS6 ratio to 16%. All P-forms (P1–P3) were sensitive to TOR inhibitors, suggesting that CrRPS6 has multiple phosphosites whose phosphorylation is dependent on TORC1. The dose-dependent effect of AZD-8055 on the phosphorylation of CrRPS6 was well suited to its effect on growth (Fig. 1C), indicating a high correlation between TORC1 activity and growth in *Chlamydomonas*.

**Fig. 1.**
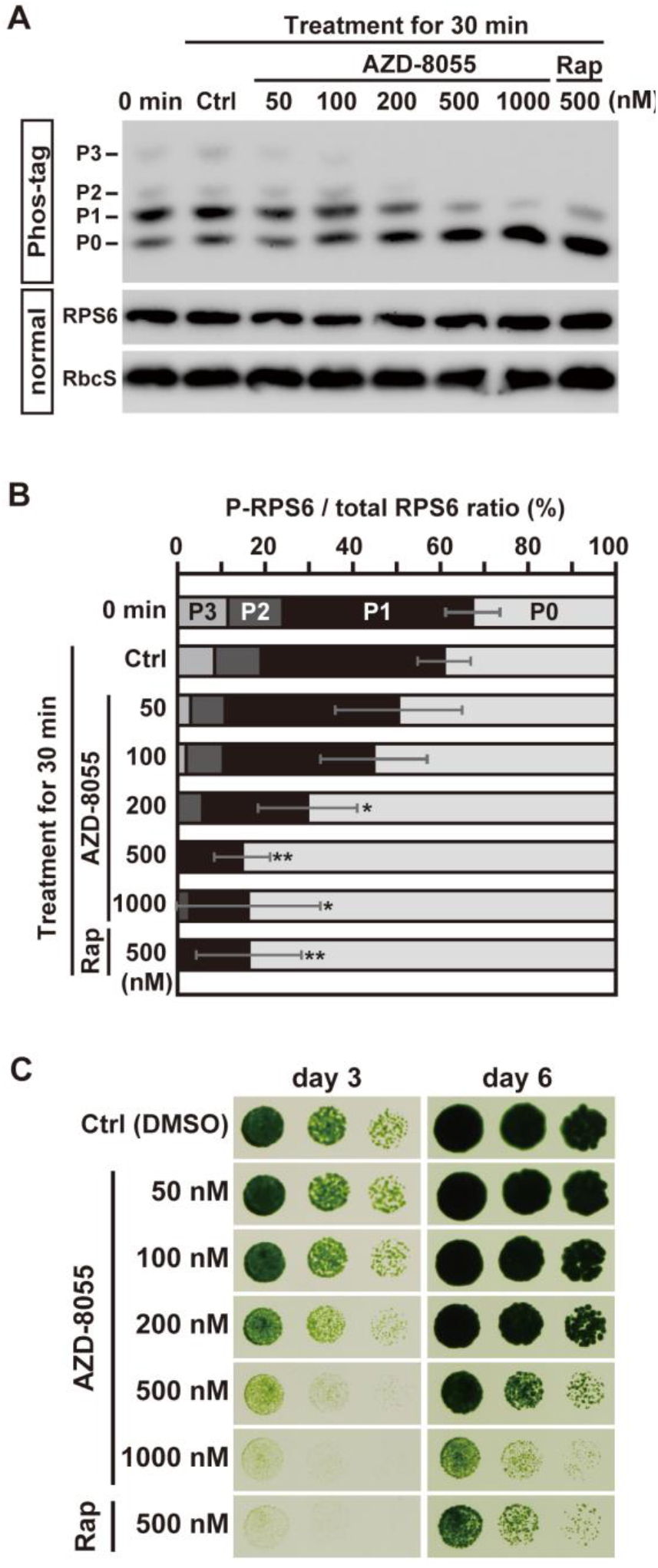
Effects of the TOR inhibitors on CrRPS6 phosphorylation status and growth of wild-type *Chlamydomonas* (strain CC-5325). **(A)** Phosphorylation status of CrRPS6 was analyzed using Phos-tag PAGE and immunoblotting using cells treated with AZD-8055 or rapamycin (Rap) for 30 min. To serve as the control (Ctrl), DMSO was added to the culture. Crude extracts (5 μg lane^−1^) were resolved via Phos-tag PAGE (Phos-tag) and normal SDS-PAGE without Phos-tag (normal), followed by immunoblotting with the antibody A (#2317S, CST). P0 in the Phos-tag gel corresponds to non-P-CrRPS6 (Fig. S2: This Fig is being generated), while P1-P3 correspond to P-forms. Rubisco small subunit (RbcS) was employed as a control. Representative data of the three biological replicates are presented. **(B)** Ratio of each form calculated from the band intensities of immunoblotting with Phos-tag PAGE analysis. Error bars (±SD) of the ratio of total P-forms (sum of P_0_ to P_1_) to total CrRPS6 are indicated. \**p* <0.05, \*\**p* <0.01 based on Student’s *t* test (vs control); *ns*, not significant. **(C)** Growth of *Chlamydomonas* on solid TAP medium with different concentrations of AZD-8055 or rapamycin. In all experiments, cells were grown under photomixotrophic conditions (TAP medium with light). Representative data of three biological replicates are shown.

### 3.2. Evaluation of TORC1 activity using the Phos-tag-based method in the diatom, *P. tricornutum*

Owing to successful application of the Phos-tag-based method to *Chlamydomonas*, this strategy was tested in ecologically important secondary lineages. The pennate diatom, *P. tricornutum*, was employed as a model. As the antibody used in *Chlamydomonas* (#2317S, CST) did not react with RPS6 in *P. tricornutum* (PtRPS6) (Fig.S3), another antibody that can react with PtRPS6 (Cat# ab225676, Abcam; denoted as Antibody B) was employed. Immunoblotting using normal SDS-PAGE resulted in a single band at 26.6 kDa (Fig. S4), which aligned with the mass obtained from the database (27.8 kDa, Phatr3_J18559 in the Ensembl database).

To determine whether the phosphorylation status of PtRPS6 was regulated by TORC1, PtRPS6 was analyzed in cells treated with various concentrations of the TOR inhibitor using Phos-tag PAGE. Due to the insensitivity of *P. tricornutum* to rapamycin [14], the ATP-competitive inhibitor, AZD-8055, was employed. Through Phos-tag PAGE and immunoblotting, at least 5 P-forms and non-P-form of PtRPS6 were detected in cells that were not treated with AZD-8055 (0 h and 0.0 μM in Fig. 2A; Fig. S2). A decrease in P-forms and a concomitant increase in non-P-forms were observed in cells treated with AZD-8055 (Fig. 2A). As quantifying individual P-forms proved difficult due to the distortion of bands (Fig. 2A), we quantified the signal intensities of whole P-PtRPS6 and non-P-PtRPS6, and then calculated the ratio of P-PtRPS6 to total PtRPS6 as an indicator of phosphorylation status (Fig. 2B). In control cells, the P-PtRPS6/total RPS6 ratio was >90% (0 h and Ctrl in Fig. 2A), which decreased to 44% with increases in the concentration of AZD-8055 up to 4 μM (Fig. 2B), indicating the major contribution of TORC1 to PtRPS6 phosphorylation.

**Fig. 2.**
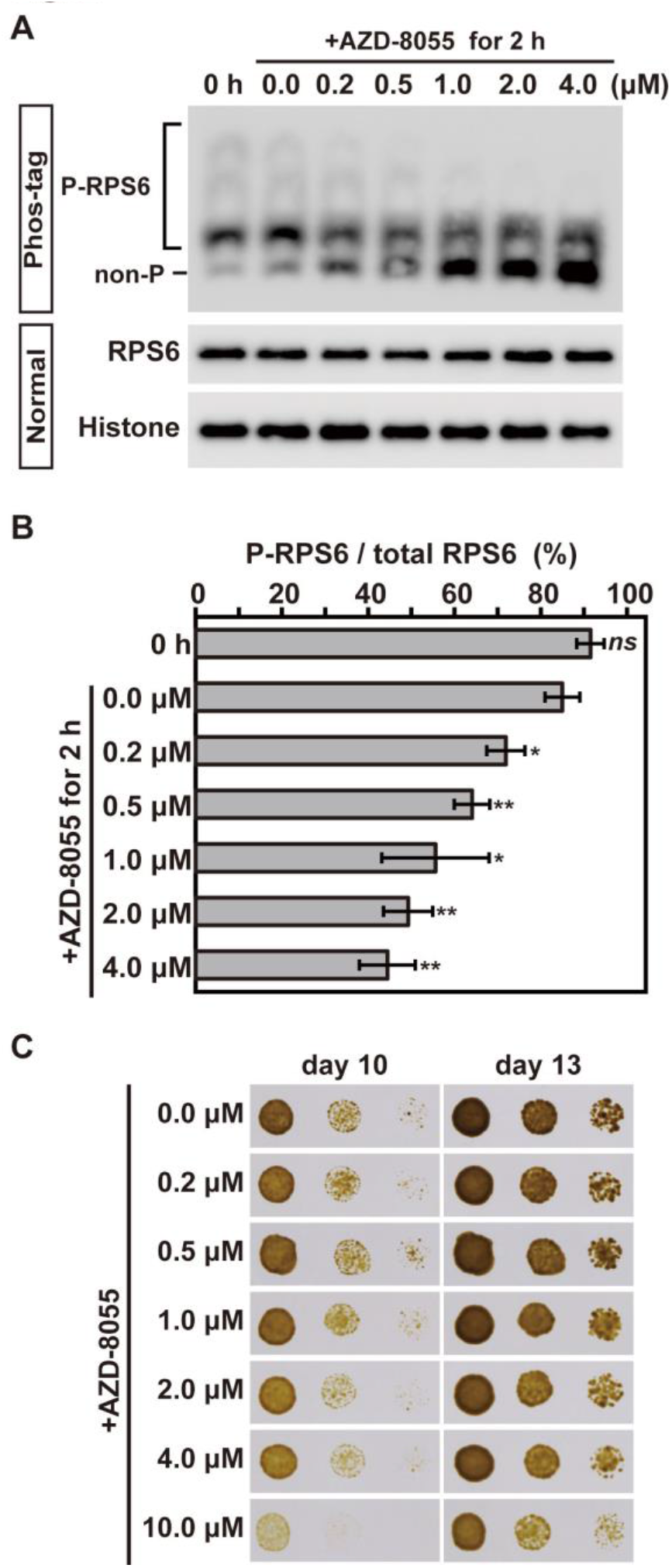
Effects of AZD-8055 on the PtRPS6 phosphorylation status and growth of wild-type *P. tricornutum*. **(A)** Phosphorylation status of PtRPS6 was analyzed using the Phos-tag-based method in cells treated with different concentrations of AZD-8055 for 2 h. In the control (Ctrl), DMSO was added to the culture. To detect PtRPS6, antibody B (#ab225676, Abcam) was used. Histone H3 was probed as a loading control. For Phos-tag PAGE (Phos-tag) and normal PAGE (normal), crude extracts containing 20 μg and 5 μg of proteins, respectively, were applied to each lane. Representative data of three biological replicates are shown. **(B)** Calculated ratio of the total P-forms to total PtRPS6. Means (±SD) of three biological replicates are shown. \**p* <0.05, \*\**p* <0.01 based on Student’s *t* test (vs control); *ns*, not significant. **(C)** Effect of AZD-8055 on the growth of *P. tricornutum* on solid MA-ESM medium. Representative data of three biological replicates are shown.

The effects of AZD-8055 on the growth of *P. tricornutum* were monitored to determine the correlation between *in vivo* TORC1 activity and growth (Fig. 2C). In contrast to *Chlamydomonas*, which displayed a good correlation between TORC1 activity and growth, no growth inhibition of *P. tricornutum* was observed with up to 1 μM of AZD-8055, despite a dose-dependent decline in the P-PtRPS6 ratio in this range (Fig. 2B&C). When the concentration of AZD-8055 was further increased, slight growth inhibition was observed with 2-4 μM, and apparent inhibition was observed with 10 μM (Fig. 2C), aligning with the results of previous studies [14,23]. As residual P-forms were still present to some extent (>40%) even at 4 μM AZD-8055 (Fig. 2B), the lack of obvious growth inhibition might be explained by the residual TORC1 activity being sufficient to maintain *P. tricornutum* growth.

### 3.3. Nutrient-dependent regulation of TORC1 in the diatom, *P. tricornutum*

The Phos-tag-based method has enabled us to monitor TORC1 activity in response to nutrient availability in *P. tricornutum*. To demonstrate TORC1 regulation by nutrients in diatoms, the time-dependent changes in the phosphorylation status of PtRPS6 under nitrogen-deficient conditions were monitored in *P. tricornutum* (Fig. 3). When cells were transferred to a nitrate-replete medium to serve as a control, the ratio of P-PtRPS6 was 90% and remained constant for 9 h (Fig. 3A and 3 B). In contrast, the P-RPS6 ratio in nitrogen-deficient medium was nearly 70% for 0 to 3 h but decreased to 13% at 9 h, indicating the inactivation of TORC1 upon the nitrogen deficiency. The chlorophyll content, which is a commonly used indicator of cellular nitrogen status, was also measured in each culture (Fig. 3C). Unlike the phosphorylation state of PtRPS6, no difference in chlorophyll content was observed between +N and -N conditions at 0 h (Fig. 3C); however, a significant increase in chlorophyll content was observed at 3 h and later in nitrogen replete cultures compared to that in nitrogen deficient cultures. The faster response of PtRPS6 phosphorylation relative to chlorophyll decline suggests the role of TORC1 as a master regulator of nitrogen deficient stress responses.

**Fig. 3.**
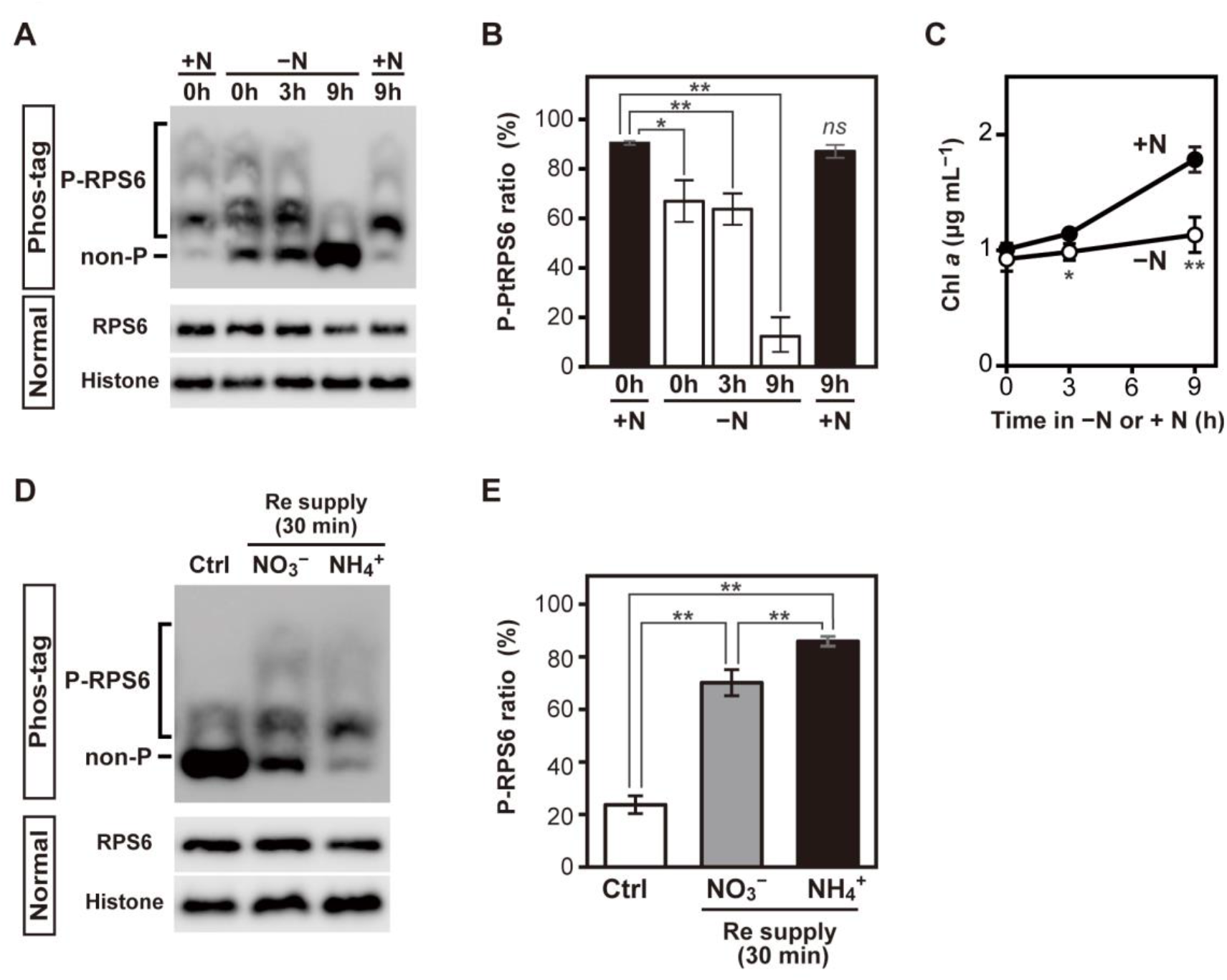
Nutrient-dependent regulation of the phosphorylation status of PtRPS6 in the diatom, *P. tricornutum*. **(A)** Changes in the phosphorylation status of PtRPS6 during nitrogen-deficient conditions based on Phos-tag PAGE and immunoblotting. **(B)** Quantified ratio of P-PtRPS6. Means (±SD) of three biological replicates are shown. \**p* <0.05, \*\**p* <0.01 based on Student’s *t* test; *ns*, not significant. **(C)** Chlorophyll contents in the nitrogen-replete and nitrogen-deficient culture. \**p* <0.05, \*\**p* <0.01 based on Student’s *t* test (+N vs −N) **(D)** Response of the phosphorylation status of PtRPS6 against resupply of nitrogen sources (NO_3_^−^, NH_4_^+^) to nitrate-deficient cells. First, cells were cultured with nitrate-free MA-ESM medium for 9 h. Subsequently, 500 μM of nitrogen sources (nitrate or ammonium) dissolved in 10 mM Tris-HCl buffer (pH8.2) was added. A blank solution (10 mM Tris-HCl, pH8.2) was employed as a control. **(E)** Quantified data of the resupply experiments. Means (±SD) of three biological replicates are shown. \*\**p* <0.01 based on Student’s *t* test.

To further demonstrate nitrogen-dependent TORC1 regulation in *P. tricornutum*, different nitrogen sources, nitrate (500 μM) or ammonium (500 μM), were added to the cells exposed to nitrogen-deficient conditions for 9 h (Fig. 3D&E). After 30 min of resupply, the P-RPS6/total RPS6 ratio was restored to 70% and 88% by nitrate (500 μM) and ammonium (500 μM), respectively; however, the ratio remained low (24%) with no nitrogen addition (Fig. 3D and E). These data indicate that TORC1 activity is regulated by the availability of N sources in *P. tricornutum*.

### 3.4. Potential of the Phos-tag-based method in diverse secondary lineages

In principle, the Phos-tag-based method can be expanded to other algae if an antibody that reacts with RPS6 in the target species is available. To broaden the range of the target species, we tested the cross-reactivity of commercial antibodies (antibodies A and B) against RPS6 from diverse secondary lineages. In particular, we selected three evolutionarily, ecologically, and industrially important secondary algae (Fig. 4A): an oleaginous stramenopile *Nannochloropsis oceanica*; *Tisochrysis lutea* (Haptophyta), and *Euglena gracilis* (Euglenid). *N. oceanica* is a model stramenopiles used to study lipid metabolism because of its high capacity to accumulate TAG [24]. *T. lutea* is used as a feed for aquaculture and can accumulate long-chain ketones (alkenones) as major storage lipids [25]. *E. gracilis* has a secondary plastid of chlorophyte origin [26] and is known to produce wax esters by fermentation [27]. When the cross-reactivity of antibody A (#2317S, CST) was tested by immunoblotting with normal SDS-PAGE, RPS6 was detected in *Chlamydomonas* and *N. oceanica* but not in other algae (Fig. 4B). In contrast, antibody B (#ab225676, Abcam) displayed broad specificity for RPS6 from all secondary algae (Fig. 4B). Although the RPS6 signal from *E. gracilis* was weaker than that from the other algae, a clear signal was detected after extended exposure (Fig. 4B, long). When these algal samples were analyzed by Phos-tag PAGE with antibody B, the bands expected to represent the P-forms were resolved in *N. oceanica* and *T. lutea* (Fig. 4C). In *N. oceanica*, at least five bands were detected, similar to that in *P. tricornutum*, which might be due to their close phylogenetic positions. In contrast, only two bands, assumed to be the P-RPS6 and non-P-form, were detected in the haptophyte, *T. lutea*. These results highlight the potential application of the Phos-tag-based method for different secondary lineages. Although no bands representing P-RPS6 were detected in *E. gracilis*, further optimization of culture conditions and/or immunoblotting protocols may enable the separation of P-forms.

**Fig. 4.**
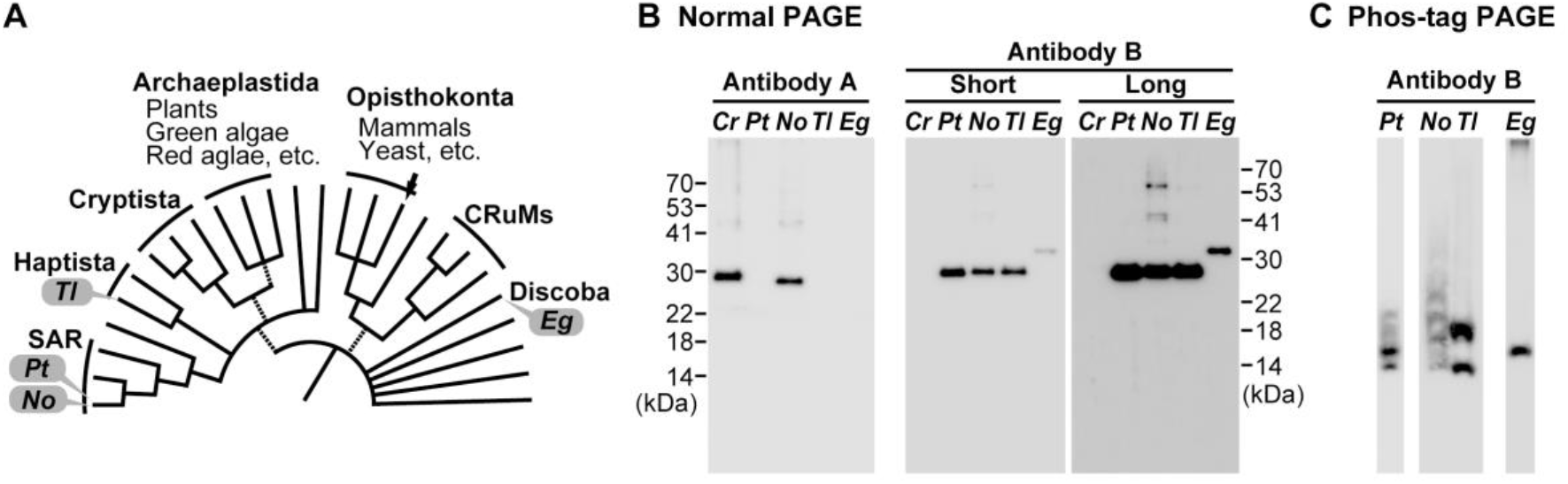
Expanding the Phos-tag-based evaluation of RPS6 phosphorylation to diverse secondary lineages. **(A)** Phylogenetic positions of secondary algae used in the experiment. Pt, *P. tricornutum*; No, *Nannochloropsis oceanica*; Tl, *Tisochrysis lutea*; Eg, *Euglena gracilis*. The tree of eukaryotes is illustrated based on Burki et al. (2020). **(B)** Testing the cross-reactivity of antibody A (#2317S, CST) and antibody B (#ab225676, Abcam) against RPS6 from diverse algae based on immunoblotting with normal SDS-PAGE. Crude extract containing 5 μg of protein was dispensed into each lane. In the immunoblot analysis with antibody B, signals obtained by short (120 s) and long (420 s) exposure are shown because the RPS6 signal in *E. gracilis* was weaker than that in other algae. **(C)** Separation and detection of putative P-RPS6 from non-P-RPS6 using Phos-tag PAGE and immunoblotting. Crude extract containing 20 μg of proteins was dispensed into each lane, except for *Chlamydomonas*, in which 5 μg was dispensed.

## 5. Discussion

Considering the conserved role of TORC1, diatom TORC1 is expected to be a key regulator of oceanic primary production under fluctuating nutrient supply. Furthermore, the “get-fat growth” induced by chemical inhibition of TORC1 in *P. tricornutum*, diatom TORC1 is the potential target of bioengineering to realize efficient biofuel production [14,15]. However, despite its biogeochemical and industrial importance, the molecular basis of TORC1 signaling in diatoms has not been explored due to the lack of methods to assess *in vivo* TORC1 activity. Therefore, we aimed to assess TORC1 activity using the RPS6 phosphorylation status as an indicator using Phos-tag PAGE and immunoblotting with a commercial antibody.

### 5.1. RPS6 phosphorylation resolved by Phos-tag PAGE in *Chlamydomonas*

Before using diatoms, we determined the feasibility of our strategy using the green alga, *Chlamydomonas.* Similar to the previous study in which P-Ser245 on CrRPS6 [9] was used as an indicator, the P-CrRPS6 ratio was found to markedly decrease to 15% in cells treated with 500 nM of rapamycin or AZD-8055 (Fig. 1), indicating the reliability of the Phos-tag based assessment of TORC1 activity. In Phos-tag PAGE, at least three P-forms of CrRPS6 were resolved (P1-P3 in Fig. 1A). In a simple interpretation, each band might represent a mono-, di-, or tri-P-form; however, caution should be exercised during interpretation because there is a case that monophosphoisotypes phosphorylated at different positions displayed different mobility shifts on Zn^2+^-Phos-tag PAGE [22]. Therefore, other possibilities remain, for instance, with P1 and P2 as mono-P-CrRPS6 that are phosphorylated at different sites, and P3 as the di-P-form. In addition to Ser245, Thr127 on CrRPS6 has been suggested to undergo TORC1-dependent phosphorylation [20]. Therefore, the three P forms may be derived from different combinations of these phosphosites, despite the potential contribution of other unidentified phosphosites.

### 5.2. Phos-tag-based method enabled monitoring of RPS6 phosphorylation status as a readout of TORC1 activity in *P. tricornutum*

Owing to its successful application in *Chlamydomonas*, we applied the same strategy to the pennate diatom, *P. tricornutum*, as a method to monitor its TORC1 activity has not been developed. A dose-dependent decrease in the P-PtRPS6 ratio following AZD-8055 treatment was also observed in *P. tricornutum* (Fig. 2), indicating the usability of the Phos-tag-based assessment of PtRPS6 phosphorylation as an indicator of TORC1 activity in this diatom. In *P. tricornutum*, more than 90% of PtRPS6 was phosphorylated in cells without inhibitor treatment (Fig. 2), which is 1.5 times higher than in *Chlamydomonas*. Due to the high basal activation state of TORC1, cells treated with up to 4 μM AZD-8055 still maintained some TORC1 activity with >40% of the P-form ratio, which might be sufficient to promote growth in the presence of AZD-8055. In addition, marine diatoms tend to exhibit tolerance to commonly used antibiotics and herbicides in saline media [28,29]. Therefore, both the high basal activation state of TORC1 and the intrinsic nature of marine diatoms might contribute to the lower sensitivity of growth to AZD-8055 in *P. tricornutum* compared to *Chlamydomonas*.

Although TORC1-dependnet phosphosites on PtRPS6 have not been identified, Phos-tag analysis revealed at least five P-forms in *P. tricornutum*, which is more than that detected in *Chlamydomonas* (three). Such variations could be derived from differences in the number of phosphosites across different organisms, as found in yeast and mammals, whose RPS6 has two and five phosphosites, respectively. Mammalian RPS6 contains regulatory phosphosites clustered at its C-terminus (Fig. S5, Ser235/236/240/244/247) and its phosphorylation occurs in an orderly manner with Ser236 as the priming site [17]. Sequence comparison revealed that three regulatory Ser residues in mammalian RPS6, including the priming site, are conserved among Stramenopiles (Fig. S5, Ser233/234/237 in *P. tricornutum*) while some diatoms including *Thalassiosira pseudonana* have Thr instead of Ser (the position of S234 in *P. tricornutum*). In addition to these shared phosphosites, a serine residue is specifically conserved in stramenopiles (Fig. S5; Ser229 in *P. tricornutum*), suggesting that stramenopiles may have shared and distinct regulatory systems compared with mammals. Despite the high conservation of the TORC1-S6K-RPS6 signaling axis among eukaryotes, little is known about the mechanisms and physiological effects of RPS6 phosphorylation in algae. Further studies using site-directed mutagenesis will provide insights into the importance of RPS6 phosphorylation in diatoms.

### 5.3. TORC1 in *P. tricornutum* is regulated by nitrogen availability

Using the Phos-tag-based method, the phosphorylation of PtRPS6 was demonstrated to markedly depend on nitrogen availability (Fig. 3), providing the first experimental evidence for the nutrient-dependent regulation of TORC1 in major oceanic primary producers. Because the removal of external nitrogen did not lead to the immediate inactivation of TORC1 activity (Fig. 3), the internal nitrogen status was suggested to be crucial for maintaining TORC1 activity during the initial phase of nitrogen starvation. Diatoms can store nitrate in vacuoles [30], which may be a source of TORC1 activity during the early phases of nitrogen starvation. The resupply of nitrogen sources to nitrogen-deficient cells quickly restored the PtRPS6 phosphorylation within 30 min (Fig. 3). Such rapid response suggests the central role of TORC1 as a master regulator of nitrogen starvation responses. Notably, the reduced form (ammonium) exhibited a more pronounced effect than the oxidized form (nitrate) (Fig. 3). Therefore, ammonium itself, or metabolites related to ammonium assimilation, such as glutamate, glutamine, or 2-oxoglutarate, might be crucial effectors for the activation of TORC1. However, as TORC1 in *A. thaliana* is activated by nitrate [3], the potential of nitrate to act as a direct effector in diatoms should not be ignored, which can be experimentally investigated using nitrate reductase mutants.

### 5.4. Wider applicability of the Phos-tag-based method in diverse secondary algae

To facilitate studies on TOR in these algae, we demonstrated the potential applicability of the Phos-tag-based method in diverse secondary lineages, such as the stramenopiles alga *N. oceanica*, haptophyte *T. lutea*, and the photosynthetic euglenid *E. gracilis* (Fig. 4). In these algae, the requisite of Phos-tag analysis (i.e., specific detection of RPS6) was achieved. In addition, Phos-tag PAGE analysis detected bands expected to be P-RPS6 in *N. oceanica* and *T. lutea*. Further optimization of the protocol may enable the evaluation of TORC1 activity in these secondary algae including *E. gracilis*. Although the regulatory system of TORC1 is well established in mammals and yeast, accumulating sequence data suggest that these findings might not be universal to eukaryotes. The Phos-tag-based method uses commercially available materials and serves as a convenient and reproducible technical platform for exploring algal TOR signaling.

## 6. Acknowledgement

The haptophyte strain *T. lutea* was kindly provided by Drs. Iwane Suzuki and Kohei Yoneda at the University of Tsukuba. *P. tricornutum*, *N. oceanica*, and *E. gracilis* were provided by the NIES through the NBRP of the MEXT. *Chlamydomonas* strain CC-5325 was obtained from Chlamydomonas Resource Center. We thank Drs. Hideya Fukuzawa and Takashi Yamano (Kyoto University) for their generous support and helpful discussions. We also thank Hiroko Shimada for technical assistance.

## 7. Author contributions

**Yoshinori Tsuji:** Conceptualization, Methodology, Validation, Investigation, Data Curation, Writing – original draft, Writing – Review & editing, Funding acquisition

**Takumi Ishikawa:** Investigation, Methodology, Validation

## 8. Funding

This work was supported by JSPS KAKENHI Grant Numbers JP20K06237 and JP23K05394 (to YT)

## 9. Declaration of competing interest

The authors declare no conflicts of interest. The authors report no commercial or proprietary interest in any product or concept discussed in this article.

## Abbreviations

MA-ESM: Marine Art SF1 enriched with modified Erd-Schreiber medium
Rag: Ras-related GTP-binding protein
Rheb: Ras homolog enriched in brain
Rop: Rho of plants
RPS6: ribosomal protein S6
S6k: ribosomal S6 kinase
TAP: tris-acetate-phosphate medium
TOR: target of rapamycin
TORC1: target of rapamycin complex 1
TORC2: target of rapamycin complex 2
λ-PPase: λ-protein phosphatase

## Notes

### Competing Interest Statement

The authors have declared no competing interest.

## References

[1] J. Simcox, D.W. Lamming, The central moTOR of metabolism, Dev. Cell 57 (2022) 691–706. 10.1016/J.DEVCEL.2022.02.024.

[2] M.J. Mallén-Ponce, M.E. Pérez-Pérez, J.L. Crespo, Deciphering the function and evolution of the target of rapamycin signaling pathway in microalgae, J. Exp. Bot. 73 (2022) 6993–7005. 10.1093/jxb/erac264.

[3] Y. Liu, X. Duan, X. Zhao, W. Ding, Y. Wang, Y. Xiong, Diverse nitrogen signals activate convergent ROP2-TOR signaling in Arabidopsis, Dev. Cell 56 (2021) 1283–1295.e5. 10.1016/j.devcel.2021.03.022.

[4] M. Schepetilnikov, J. Makarian, O. Srour, A. Geldreich, Z. Yang, J. Chicher, P. Hammann, L.A. Ryabova, GTPase ROP2 binds and promotes activation of target of rapamycin, TOR, in response to auxin, EMBO J. 36 (2017) 886–903. 10.15252/embj.201694816.

[5] P. Cao, S.-J. Kim, A. Xing, C.A. Schenck, L. Liu, N. Jiang, J. Wang, R.L. Last, F. Brandizzi, Homeostasis of branched-chain amino acids is critical for the activity of TOR signaling in Arabidopsis, Elife 8 (2019) e50747. 10.7554/eLife.50747.

[6] C. Ingargiola, I. Jéhanno, C. Forzani, A. Marmagne, J. Broutin, G. Clément, A.-S. Leprince, C. Meyer, The Arabidopsis Target of Rapamycin kinase regulates ammonium assimilation and glutamine metabolism, Plant Physiol. 192 (2023) 2943–2957. 10.1093/plphys/kiad216.

[7] I. Pancha, K. Chokshi, K. Tanaka, S. Imamura, Microalgal target of rapamycin (TOR): a central regulatory hub for growth, stress response and biomass production, Plant Cell Physiol. 61 (2020) 675–684. 10.1093/PCP/PCAA023.

[8] S. Upadhyaya, S. Agrawal, A. Gorakshakar, B.J. Rao, TOR kinase activity in *Chlamydomonas reinhardtii* is modulated by cellular metabolic states, FEBS Lett 594 (2020) 3122–3141. 10.1002/1873-3468.13888.

[9] I. Couso, M.E. Pérez-Pérez, M.M. Ford, E. Martínez-Force, L.M. Hicks, J.G. Umen, J.L. Crespo, Phosphorus availability regulates TORC1 signaling via LST8 in Chlamydomonas, Plant Cell 32 (2020) 69–80. 10.1105/tpc.19.00179.

[10] M.J. Mallén-Ponce, M.E. Pérez-Pérez, J.L. Crespo, Photosynthetic assimilation of CO_2_ regulates TOR activity, Proc. Natl. Acad. Sci. USA. 119 (2022) e2115261119. 10.1073/pnas.2115261119.

[11] H. Mulvey, L. Dolan, RHO of plant signaling was established early in streptophyte evolution, Current Biology (2023) 5515–5525. 10.1016/j.cub.2023.11.007.

[12] R.G. Dorrell, F. Liu, C. Bowler, Reconstructing dynamic evolutionary events in diatom nuclear and organelle genomes, in: The Molecular Life of Diatoms, Springer International Publishing, Cham, 2022: pp. 147–177. 10.1007/978-3-030-92499-7_6.

[13] T. Tanaka, Y. Maeda, A. Veluchamy, M. Tanaka, H. Abida, E. Maréchal, C. Bowler, M. Muto, Y. Sunaga, M. Tanaka, T. Yoshino, T. Taniguchi, Y. Fukuda, M. Nemoto, M. Matsumoto, P.S. Wong, S. Aburatani, W. Fujibuchi, Oil accumulation by the oleaginous diatom *Fistulifera solaris* as revealed by the genome and transcriptome, Plant Cell 27 (2015) 162–76. 10.1105/tpc.114.135194.

[14] L. Prioretti, L. Avilan, F. Carrière, M.-H. Montané, B. Field, G. Grégori, B. Menand, B. Gontero, The inhibition of TOR in the model diatom *Phaeodactylum tricornutum* promotes a get-fat growth regime, Algal Res. 26 (2017) 265–274. 10.1016/j.algal.2017.08.009.

[15] L. Prioretti, F. Carriere, B. Field, L. Avilan, M.-H. Montané, B. Menand, B. Gontero, Targeting TOR signaling for enhanced lipid productivity in algae, Biochimie 169 (2020) 12–17. 10.1016/j.biochi.2019.06.016.

[16] A. González, M. Shimobayashi, T. Eisenberg, D.A. Merle, T. Pendl, M.N. Hall, T. Moustafa, TORC1 promotes phosphorylation of ribosomal protein S6 via the AGC kinase Ypk3 in *Saccharomyces cerevisiae*, PLoS One 10 (2015) e0120250. 10.1371/journal.pone.0120250.

[17] O. Meyuhas, Ribosomal protein S6 phosphorylation: four decades of research., Int. Rev. Cell Mol. Biol. 320 (2015) 41–73. 10.1016/bs.ircmb.2015.07.006.

[18] Y. Xiong, M. McCormack, L. Li, Q. Hall, C. Xiang, J. Sheen, Glucose–TOR signalling reprograms the transcriptome and activates meristems, Nature 496 (2013) 181–186. 10.1038/nature12030.

[19] I. Couso, B.S. Evans, J. Li, Y. Liu, F. Ma, S. Diamond, D.K. Allen, J.G. Umen, Synergism between inositol polyphosphates and TOR kinase signaling in nutrient sensing, growth control, and lipid metabolism in *Chlamydomonas*, Plant Cell 28 (2016) 2026–2042. 10.1105/tpc.16.00351.

[20] E.G. Werth, E.W. McConnell, I. Couso Lianez, Z. Perrine, J.L. Crespo, J.G. Umen, L.M. Hicks, Investigating the effect of target of rapamycin kinase inhibition on the *Chlamydomonas reinhardtii* phosphoproteome: from known homologs to new targets, New Phytol. 221 (2019) 247–260. 10.1111/nph.15339.

[21] A. Danbara, Y. Shiraiwa, The requirement of selenium for the growth of marine Coccolithophorids, *Emiliania huxleyi*, *Gephyrocapsa oceanica* and *Helladosphaera* sp. (Prymnesiophyceae), Plant Cell Physiol. 40 (1999) 762–766. 10.1093/oxfordjournals.pcp.a029603

[22] E. Kinoshita, E. Kinoshita-Kikuta, Improved Phos-tag SDS-PAGE under neutral pH conditions for advanced protein phosphorylation profiling, Proteomics 11 (2011) 319–323. 10.1002/pmic.201000472.

[23] D.Y. Kwon, T.T. Vuong, J. Choi, T.S. Lee, J.-I. Um, S.Y. Koo, K.T. Hwang, S.M. Kim, Fucoxanthin biosynthesis has a positive correlation with the specific growth rate in the culture of microalga *Phaeodactylum tricornutum*, J. Appl. Phycol. 33 (2021) 1473–1485. 10.1007/s10811-021-02376-5.

[24] T. Nobusawa, K. Hori, H. Mori, K. Kurokawa, H. Ohta, Differently localized lysophosphatidic acid acyltransferases crucial for triacylglycerol biosynthesis in the oleaginous alga *Nannochloropsis*, Plant J. 90 (2017) 547–559. 10.1111/tpj.13512.

[25] Y. Tsuji, M. Yoshida, Biology of haptophytes: complicated cellular processes driving the global carbon cycle, in: Y. Hirakawa (Ed.), Adv. Bot. Res., 2017: pp. 219–261. 10.1016/bs.abr.2017.07.002.

[26] K.E. Wiegert, M.S. Bennett, R.E. Triemer, Evolution of the chloroplast genome in photosynthetic euglenoids: a comparison of *Eutreptia viridis* and *Euglena gracilis* (Euglenophyta), Protist 163 (2012) 832–843. 10.1016/j.protis.2012.01.002.

[27] T. Ogawa, M. Tamoi, A. Kimura, A. Mine, H. Sakuyama, E. Yoshida, T. Maruta, K. Suzuki, T. Ishikawa, S. Shigeoka, Enhancement of photosynthetic capacity in *Euglena gracilis* by expression of cyanobacterial fructose-1,6-/sedoheptulose-1,7-bisphosphatase leads to increases in biomass and wax ester production, Biotechnol. Biofuels 8 (2015) 80. 10.1186/s13068-015-0264-5.

[28] P. Kroth, Molecular Biology and the Biotechnological Potential of Diatoms, in: R. León, A. Galván, E. Fernández (Eds.), Transgenic microalgae as green cell factories, Springer New York, New York, NY, 2007: pp. 23–33. 10.1007/978-0-387-75532-8_3.

[29] L.A. Zaslavskaia, J.C. Lippmeier, P.G. Kroth, A.R. Grossman, K.E. Apt, Transformation of the diatom *Phaeodactylum tricornutum* (Bacillariophyceae) with a variety of selectable marker and reporter genes, J Phycol 36 (2001) 379–386. 10.1046/j.1529-8817.2000.99164.x.

[30] J.K. McCarthy, S.R. Smith, J.P. McCrow, M. Tan, H. Zheng, K. Beeri, R. Roth, C. Lichtle, U. Goodenough, C.P. Bowler, C.L. Dupont, A.E. Allen, Nitrate reductase knockout uncouples nitrate transport from nitrate assimilation and drives repartitioning of carbon flux in a model pennate diatom, Plant Cell 29 (2017) 2047–2070. 10.1105/tpc.16.00910.

