## Supplementary material for "Assessing Target of Rapamycin (TOR) activity in the diatom *Phaeodactylum tricornutum* using commercially available materials": Fig. S

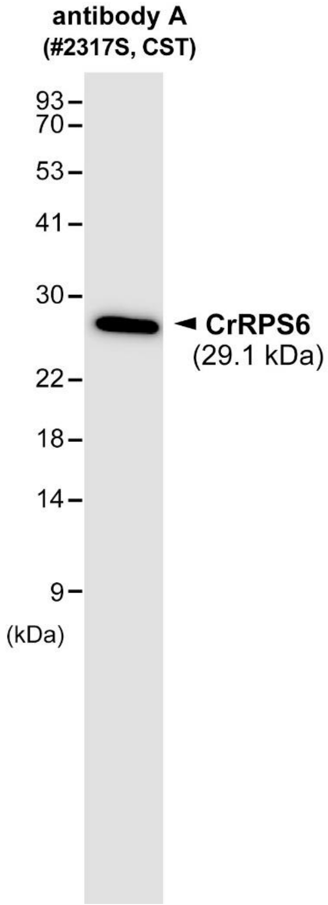


**Fig. S1** Detection of *Chlamydomonas* RPS6 (CrRPS6) by immunoblotting with a commercial antibody (antibody A, Cell Signaling Technology, #2317S). Crude extracts containing 5 μg of proteins lane$^{-1}$ were resolved by normal SDS-PAGE (w/o Phos-tag), followed by immunoblotting with antibody A.

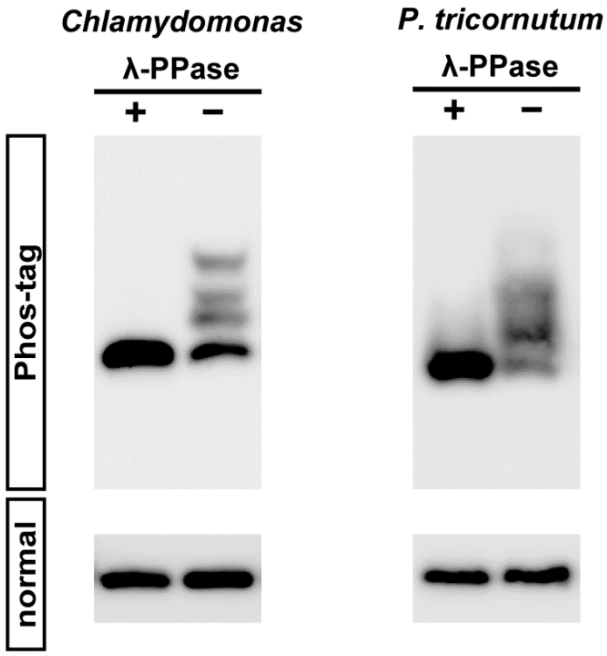


**Fig. S2** Determination of a band corresponding to dephosphorylated RPS6 by λ-PPase treatment. Crude extracts treated with λ-PPase (+) and non-treated samples were analyzed by Phos-tag PAGE and immunoblotting (upper) and normal PAGE (lower).

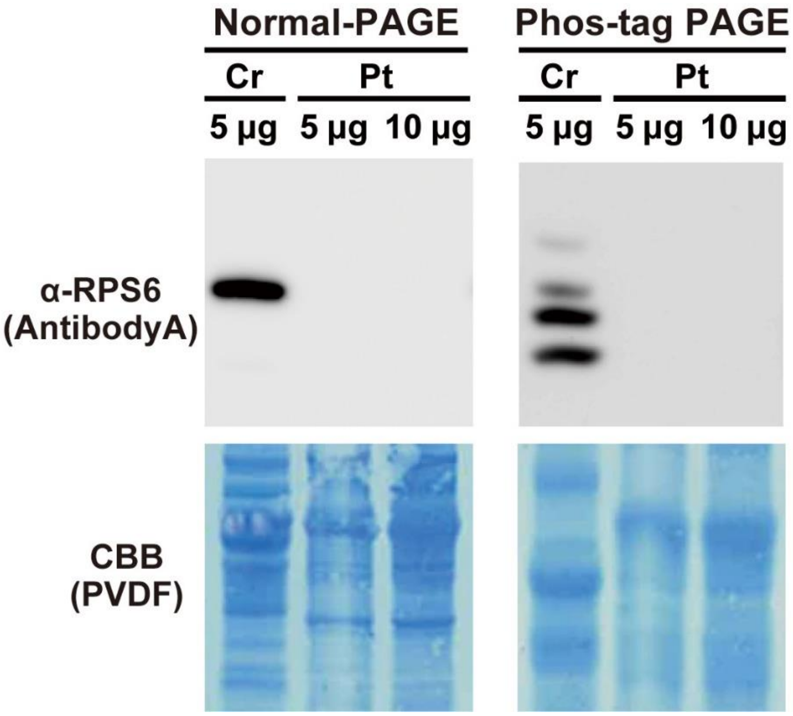


**Fig. S3** Antibody A did not cross-react with RPS6 in *P. tricornutum*. Crude extracts prepared from *Chlamydomonas* (Cr) or *P. tricornutum* (Pt) were analyzed by normal SDS-PAGE (w/o Phos-tag) and Phos-tag PAGE (w/ Phos-tag). The amount of crude proteins loaded on each lane was indicated.

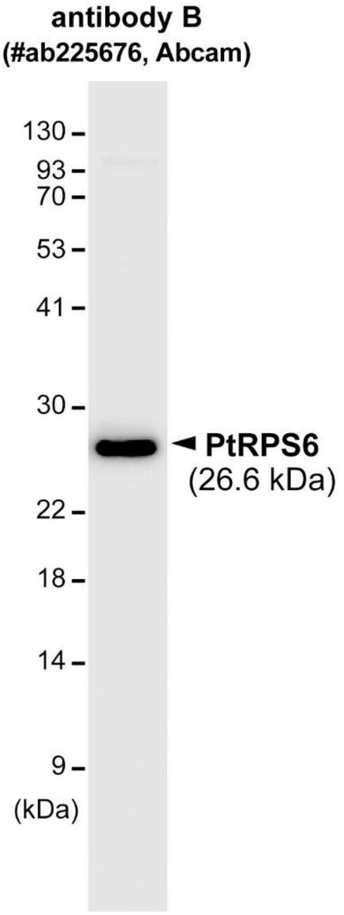


**Fig. S4** Detection of *P. tricornutum* RPS6 (PtRPS6) by immunoblotting with a commercial antibody (antibody B, Abcam, #225676). Crude extracts containing 5 μg of proteins lane$^{-1}$ were resolved by normal SDS-PAGE (w/o Phos-tag), followed by immunoblotting with antibody B.

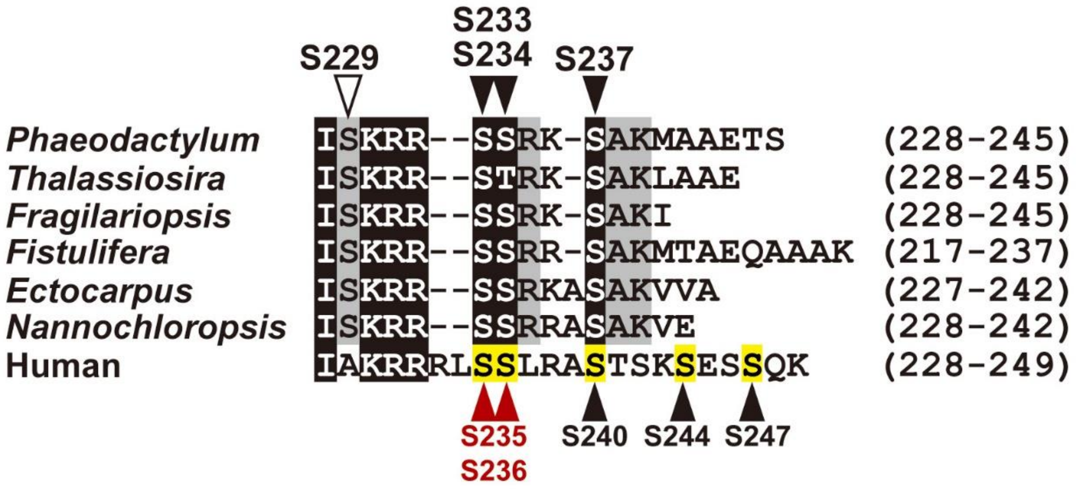


**Fig. S5** Comparison of C-terminus regulatory region in RPS6 from stramenopiles and human. Residues conserved in stramenopiles or all organisms are indicated by black or gray, respectively. The TORC1-dependent phosphosites in human RPS6 were highlighted in yellow. Red allows (S235/236) indicate the priming sites of human RPS6. The Ser residue specifically conserved in stramenopiles is shown by the white arrow. Sequences used are; Uniprot A2A3R6 (human RPS6), Genbank EED88801 (*Thalassiosira pseudonana*), Genbank OEU21702 (*Fragilariopsis cylindrus*), Genbank GAX12748 (*Fistulifera solaris*), Genbank CBJ34010 (*Ectocarpus siliculosus*), Genbank EWM21878 (*Nannochloropsis gaditana*), Uniprot A2A3R6 (human).
